# Dual targeting of TatA points to a chloroplast-like Tat pathway in plant mitochondria

**DOI:** 10.1101/2020.04.06.026997

**Authors:** Bationa Bennewitz, Mayank Sharma, Franzisca Tannert, Ralf Bernd Klösgen

## Abstract

The biogenesis of membrane-bound electron transport chains requires membrane translocation pathways for folded proteins carrying complex cofactors, like the Rieske Fe/S proteins. Two independent systems were developed during evolution, namely the Twin-arginine translocation (Tat) pathway, which is present in bacteria and chloroplasts, and the Bcs1 pathway found in mitochondria of yeast and mammals. Mitochondria of plants carry a Tat-like pathway which was hypothesized to operate with only two subunits, a TatB-like protein and a TatC homolog (OrfX), but lacking TatA. Here we show that the nuclearly encoded TatA has dual targeting properties, i.e., it can be imported into both, chloroplasts and mitochondria. Dual targeting of TatA was observed with *in organello* experiments employing isolated chloroplasts and mitochondria as well as after transient expression of suitable reporter constructs in epidermal leaf cells. The extent of transport of these constructs into mitochondria of transiently transformed leaf cells was relatively low, causing a demand for highly sensitive methods to be detected, like the *sa*splitGFP approach. Yet, the dual import of TatA into mitochondria and chloroplasts observed here points to a common mechanism of Tat transport for folded proteins within both endosymbiotic organelles.

## Introduction

Transport of folded proteins across membranes is a major challenge in a cell. While unfolded polypeptides have a largely constant diameter and thus require a membrane pore of essentially constant size, the different sizes of folded passenger proteins demand for sophisticated transport machineries and mechanisms. This is particularly demanding in membrane systems comprising an electron transport chain, like the inner mitochondrial membrane, the thylakoid membrane, and the cytoplasmic membrane of bacteria (Berry, 2003). In these membranes, a membrane potential required for ATP synthesis is generated which has to be maintained also during transport of folded proteins. On the other hand, the functionality of the electron transport chains depends also on correctly assembled cofactors. Some complex cofactors are transported across the membranes only after incorporation into the respective apoproteins. In consequence, these proteins must be at least partially folded prior to the actual membrane translocation. Typical examples are the Rieske Fe/S proteins found in respiratory and photosynthetic electron transport chains. These proteins get their Fe/S-clusters incorporated on the *cis*-side of the respective membrane, i.e. in the bacterial cytoplasm, chloroplast stroma, or mitochondrial matrix (Lill and Mühlenhoff, 2008; Balk and Schaedler, 2014; Blanc et al., 2015), while they exert their activity on the *trans*-side of the membrane, i.e., the bacterial periplasmic space, the thylakoid lumen, or the mitochondrial inner membrane space, respectively.

In the cytoplasmic membrane of bacteria and the thylakoid membrane of chloroplasts, transport of such folded proteins is executed by the Twin-arginine translocation (Tat) machinery which consists of three membrane integral subunits named TatA, TatB, and TatC (in the thylakoid system also addressed as Tha4, Hcf106, and cpTatC, respectively; Müller and Klösgen, 2005). In contrast, in yeast mitochondria a single membrane protein, Bcs1, was found to be necessary and sufficient for the transport of the folded Rieske Fe/S protein (Wagener et al., 2011). However, this might not hold true for all mitochondria because the Bcs1-protein of plants is N-terminally truncated and thus presumably not able to provide this function (Carrie et al., 2016). Instead, plant mitochondria apparently house a Tat-like pathway, sínce a protein with homology to TatC, OrfX, was found to be encoded in the mitochondrial genome (Sünkel et al., 1994; Braun and Schmitz, 1999). Moreover, a TatB-like protein was recently identified which shows considerable structural homology to TatB (Carrie et al., 2016), although sequence analyses suggest that the two genes are not derived from a common ancestor. Recently, it was shown that this TatB-like protein is in fact essential for the topogenesis of plant mitochondrial Rieske proteins (Schäfer et al., 2020).

Yet, a mitochondrial TatA protein has not yet been identified to date, neither by homology searches nor by structure-derived analyses. This led to the suggestion that the Tat machinery of plant mitochondria consists of only two subunits, namely the TatB-like protein and mitochondrially encoded TatC, in analogy to the Tat machineries of Gram-positive bacteria like *Bacillus subtilis* which consist of TatA and TatC only (Petru et al., 2018). A possible alternative to the lack of TatA would be that the protein is instead dually targeted, i.e. transported not only into chloroplasts but also into mitochondria, as was found for approx. 5% of the nuclear encoded organelle proteins (Mitschke et al., 2009; Baudisch et al., 2014). Here, we have applied different approaches, *in vivo* as well as *in organello*, to examine this possibility.

## Results

### TatA shows differential organelle targeting in transiently transformed plant cells

As the first step to investigate the organelle targeting specificity of TatA, epidermal cells of pea leaves were transiently transformed by biolistic transformation with a gene encoding a chimeric reporter protein under the control of the constitutive 35S promoter of Cauliflower Mosaic Virus. The reporter consisted of the N-terminal 100 residues of pea TatA comprising the organelle targeting transit peptide fused to the *enhanced Yellow Fluorescent Protein* (eYFP). Subcellular localization of eYFP within the transformed cells was analyzed by confocal laser scanning microscopy. As expected, all transformed cells showed localization of the eYFP reporter in chloroplasts (Fig. 1A), in line with the fact that authentic TatA is a component of the thylakoidal Twin-arginine protein transport (Tat) machinery (Müller and Klösgen, 2005). In a number of cells though, eYFP fluorescence could additionally be detected also in highly mobile punctuate structures strongly resembling mitochondria (e.g., Fig. 1A, panel I). In fact, expression of a comparable gene construct comprising the N-terminal 100 residues of the mitochondrial Rieske protein fused to eYFP showed the same punctuate fluorescence pattern (Fig. 1B) confirming that these structures are indeed mitochondria. If instead the transport signal of a strictly chloroplast-specific protein like TatB was used for targeting of the eYFP reporter, no such structures in addition to chloroplasts could be detected (Fig. 1C).

**Figure 1.**
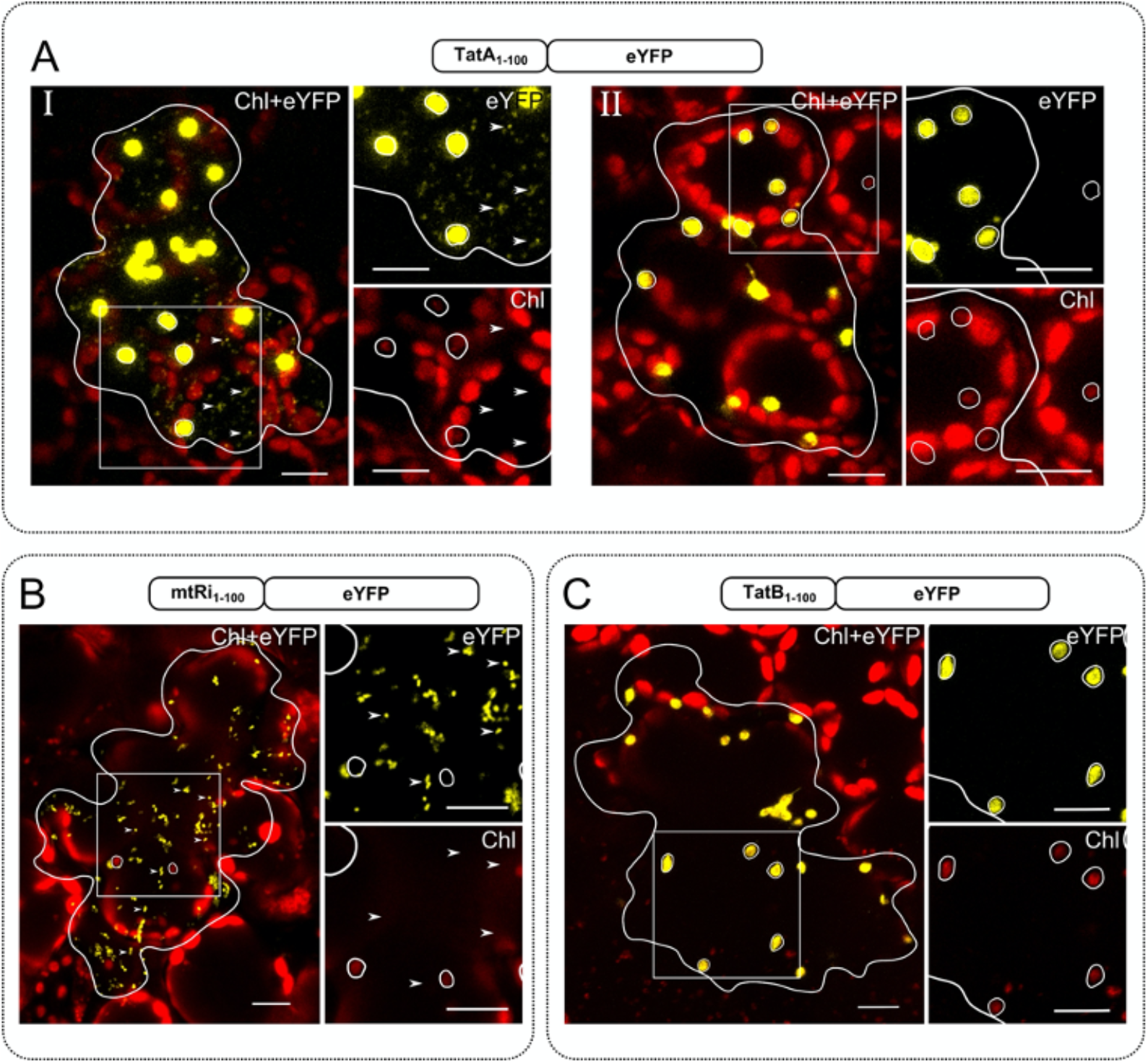
Localization of the TatA_1-100_/eYFP chimera in pea. **(A)** The coding sequence of peaTatA_1-100_/eYFP was transiently expressed under the control of the CaMV 35S promoter after particle bombardment of leaf epidermis cells of pea and analyzed by confocal laser scanning microscopy. Representative cells showing either dual localization of the candidate proteins in both mitochondria and chloroplasts (panel I) or solely in chloroplasts (panel II) are presented as overlay images of the chlorophyll channel (displayed in *red*) and the eYFP channel (displayed in *yellow*). Localization of mitochondrial (mtRi_1-100_/eYFP) and chloroplast control proteins (TatB_1-100_/eYFP) analyzed in parallel is shown in **(B)** and **(C)**, respectively. The borders of the transformed cells are depicted by a continuous *white line*. The strong chlorophyll signals in the background are derived from the larger chloroplasts of untransformed mesophyll cells underneath the epidermal cell layers. The *squares* highlight areas of the transformed cells that are shown in higher magnification separately for the chlorophyll channel and the eYFP channel, as indicated. The position of representative plastids of each transformed cell is encircled for better visualization. Representative mitochondria are marked with an *arrowhead*. The scale bars correspond to 10 μm.

The signal strength of eYFP fluorescence in the mitochondria of cells expressing the TatA_1-100_/eYFP construct was usually quite low. Furthermore, in all repetitions of the experiment only approximately 5 - 15 % of the transformed cells showed dual targeting of the reporter also into mitochondria, while in the majority of cells solely import into chloroplasts could be detected. Remarkably, even cells in close proximity to each other can show deviating organelle targeting specificity (suppl. Fig. S1), which rules out that the differences in the targeting specificity observed result from variation of the transformation conditions.

Since different transformation systems sometimes lead to deviating targeting results (Sharma et al., 2018), we used a second approach to determine the targeting properties of TatA, notably agroinfiltration into *Nicotiana benthamiana.* In this case, epidermal cells from *Nicotiana* leaf tissue were transiently transformed with the peaTatA_1-100_/eYFP gene construct described above leading to a large number of transformed cells at the same time. Unexpectedly, all transformed cells showed localization of the eYFP reporter solely within chloroplasts, while fluorescence in mitochondria could not be detected (Fig. 2A). In contrast, expression of the control construct mtRi_1-100_/eYFP showed clear fluorescence signals in mitochondria (Fig. 2B) which rules out that mitochondrial fluorescence could on principle not be detected in these assays.

**Figure 2.**
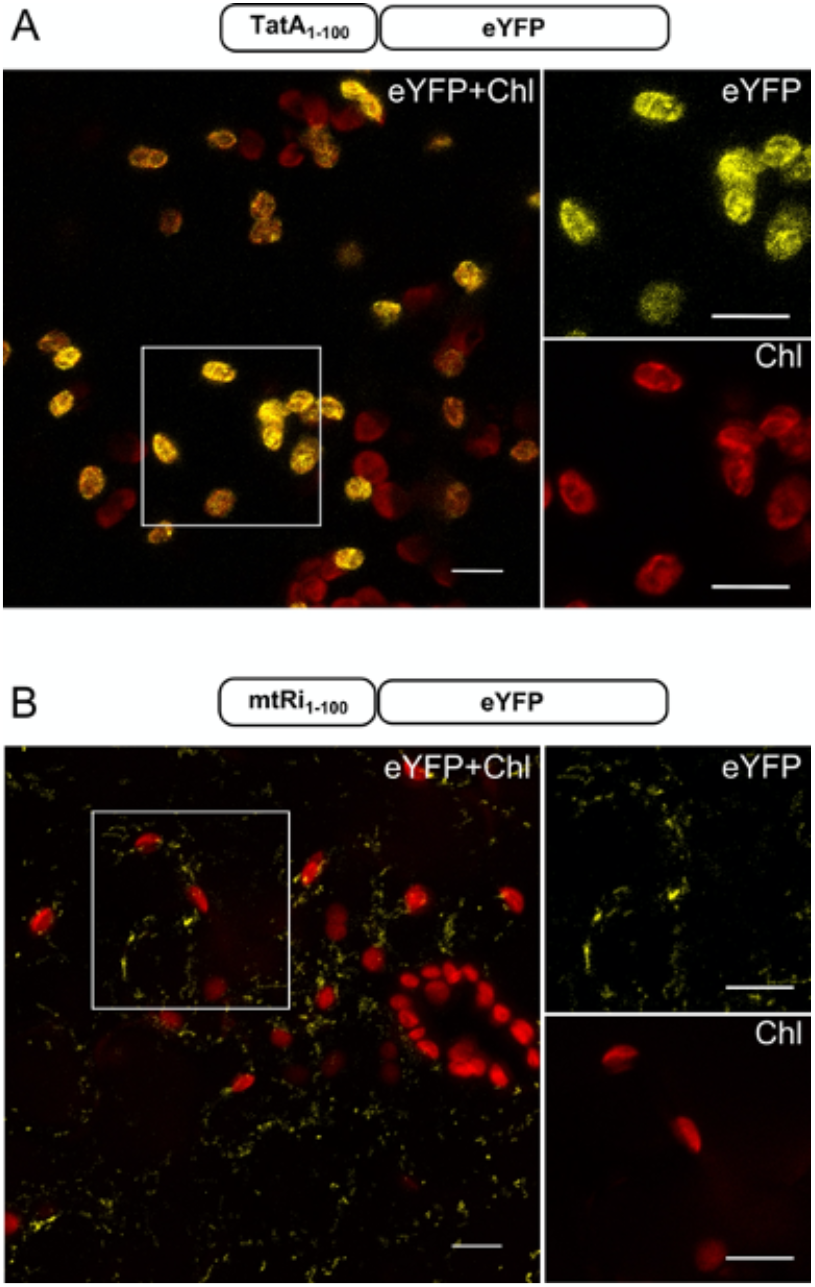
Monospecific targeting of TatA_1-100_/eYFP in *Nicotiana benthamiana* after *Agrobacterium* infiltration. Confocal laser scanning microscopy of the lower epidermis of *Nicotiana benthamiana* infiltrated with *Agrobacterium tumefaciens* strain GV3101 carrying constructs encoding TatA_1-100_/eYFP **(A)** or mtRi_1-100_/eYFP **(B)**. For further details see the legend of Fig. 1. Scale bars: 10 μm.

This result suggests at first glance that in *Nicotiana benthamiana* transport of peaTatA_1-100_/eYFP into mitochondria does not take place. However, considering the low number of cells showing dual targeting of the TatA_1-100_/eYFP reporter construct after biolistic transformation and the weak fluorescence signals observed in mitochondria in these assays (Fig. 1A), it might also be possible that mitochondrial transport of the reporter in *Nicotiana* epidermal cells was not sufficiently strong to be detected. In particular, since the large number of surrounding cells showing strong fluorescence signals in chloroplasts might overlay faint signals in mitochondria.

### Agroinfiltration with self-assembling GFP allows for more sensitive detection of organelle targeting

In order to circumvent such sensitivity problems, the approach was therefore modified making use of the self-assembling split green fluorescent protein (*sa*split-GFP) technology (Cabantous et al., 2005) which rests on the self-assembly properties of two fragments of an engineered ‘superfolder’ GFP variant, sfGFP. The “receptor” fragment GFP1-10 comprises the ten N-terminal antiparallel β-sheets of sfGFP, while GFP11 represents the C-terminal eleventh β-sheet. The two non-fluorescent fragments can assemble spontaneously to reconstitute the functional fluorophore. If combined with specific organelle-targeting signals, the self-assembly property can be used for subcellular localization studies (Sharma et al. 2019; see also suppl. Fig. S2).

The subcellular localization of TatA was analyzed by combining the N-terminal 100 residues of peaTatA with a sevenfold repeat of GFP11 (GFP11_x7_) (Sharma et al., 2019). The resulting chimera (peaTatA_1-100_/GFP11_x7_) was co-infiltrated with the organelle-specific constructs FNR_1-55_/GFP1-10 or mtRi_1-100_/GFP1-10, which mediate transport of the “receptor” component specifically into chloroplasts and mitochondria, respectively. With this approach, it was indeed possible to detect transport of the TatA_1-100_/GFP11_x7_ construct not only into the chloroplasts of *Nicotiana* cells but also into small, highly mobile punctuate structures (Fig. 3) which were confirmed as mitochondria by staining with MitoTracker Orange (Poot et al., 1996). The fluorescence signal intensity within mitochondria was again clearly weaker than in chloroplasts, in accordance with the results after biolistic transformation (Fig. 1A). This indicates that the extent of transport into mitochondria mediated by the TatA transit peptide is relatively low which might explain why mitochondrial transport was not detected by agroinfiltration with the peaTatA_1-100_/EYFP-construct (Fig. 2).

**Figure 3.**
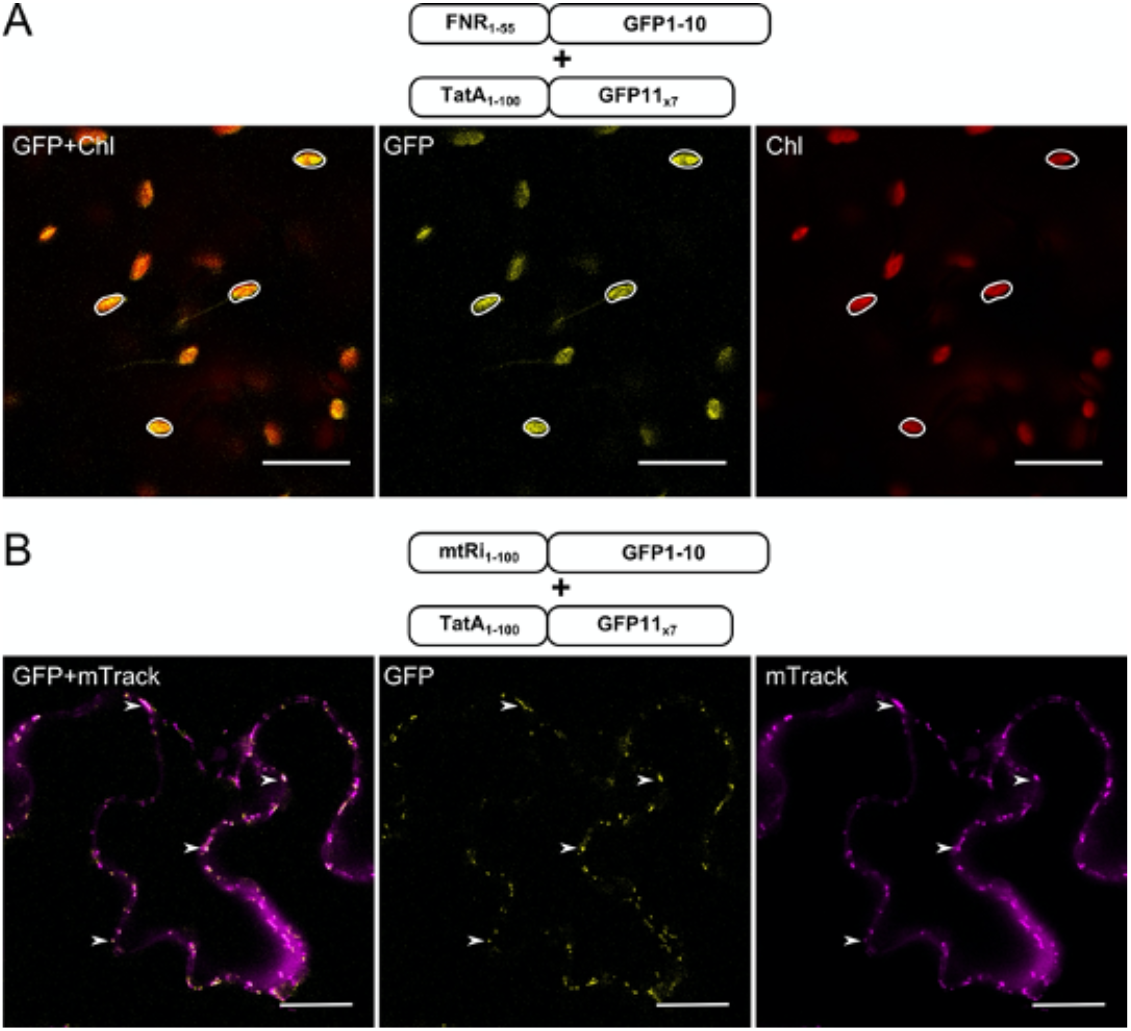
Dual targeting of TatA_1-100_ with the sasplit-GFP approach. The coding sequence of TatA_1-100_/GFP11_x7_) was transiently expressed with either FNR1-55/GFP1-10 **(A)** or mtRi_1-100_/GFP1-10 **(B)** via *Agrobacterium* co-infiltration into the lower epidermis of *Nicotiana benthamiana* leaves and analyzed by confocal laser scanning microscopy. Representative images, which show targeting of the candidate protein into chloroplasts **(A),** are presented in the chlorophyll channel (right panel, displayed in *red*), the GFP channel (middle panel, displayed in *green*) and as overlay image (left panel). Targeting of the candidate protein into mitochondria **(B)** is shown with MitoTracker Orange (right panel, displayed in *magenta*), in the GFP channel (middle panel) and as overlay image (left panel). For further details see the legend of Fig. 1. Scale bars: 10 μm.

Remarkably, the number of cells showing fluorescence in chloroplast vs. mitochondria in these experiments was considerably different from each other. While GFP reconstitution within chloroplasts was always observed in a large number of cells after transformation, fluorescence signals in mitochondria could be detected with only low frequency in all repetitions. This is not a matter of lacking reconstitution in mitochondria in general because GFP fluorescence was observed at high frequency with the mtRi_1-100_/GFP11_x7_ construct (Sharma et al. 2019; see also suppl. Fig. S2). Unfortunately, it is on principle not possible with such assays to quantify the exact ratio of organellar targeting per number of transformed cells, since transformed cells without co-localization of GFP1-10 and GFP11 cannot be distinguished from untransformed cells.

### TatA is imported into isolated intact mitochondria and chloroplasts

The apparent variability in the results regarding the mitochondrial targeting properties of the peaTatA transit peptide demanded for further examination. We have chosen *in organello* protein transport experiments for this purpose which rest on the analysis of authentic precursor proteins rather than chimeric reporter constructs. The precursor proteins are obtained in radiolabelled form by *in vitro* transcription/translation in the presence of 35S-methionine. Incubation of such *in vitro* generated TatA precursor protein with either chloroplasts or mitochondria isolated from pea leaves yielded in both instances a processing product of similar size (approx. 15 kDa, Fig. 4). This value deviates significantly from the molecular weight of mature TatA as deduced from the corresponding cDNA sequence (8.9 kDa). Such aberrant mobility on SDS-PAGE was observed already earlier for TatA (Jakob et al., 2009). The processing products are resistant against protease added externally to the assays after transport which demonstrates that the proteins were not merely bound to the organellar surfaces but have in fact been internalized. In accordance with the results of the *in vivo* experiments shown above, the amount of mature TatA accumulating in the chloroplasts is apparently higher than in mitochondria.

Control precursor proteins for organelle transport, which were analyzed in parallel, showed the expected monospecific organelle transport into either chloroplasts (TatB) or mitochondria (mtRieske) (Fig. 4). GrpE, on the other hand, which had been shown already earlier to possess dual targeting properties (Baudisch et al., 2014), is imported into both organelles, as expected. In this case, the processing products accumulating in chloroplasts and mitochondria are different from each other (Fig. 4) demonstrating that the processing peptidases of chloroplasts and mitochondria (SPP and MPP, respectively) recognize different cleavage sites within the GrpE precursor. These different processing products allow furthermore to estimate the degree of cross-contaminating organelles because small amounts of terminally processed mitochondrial GrpE are found also in chloroplasts and *vice versa*.

**Figure 4.**
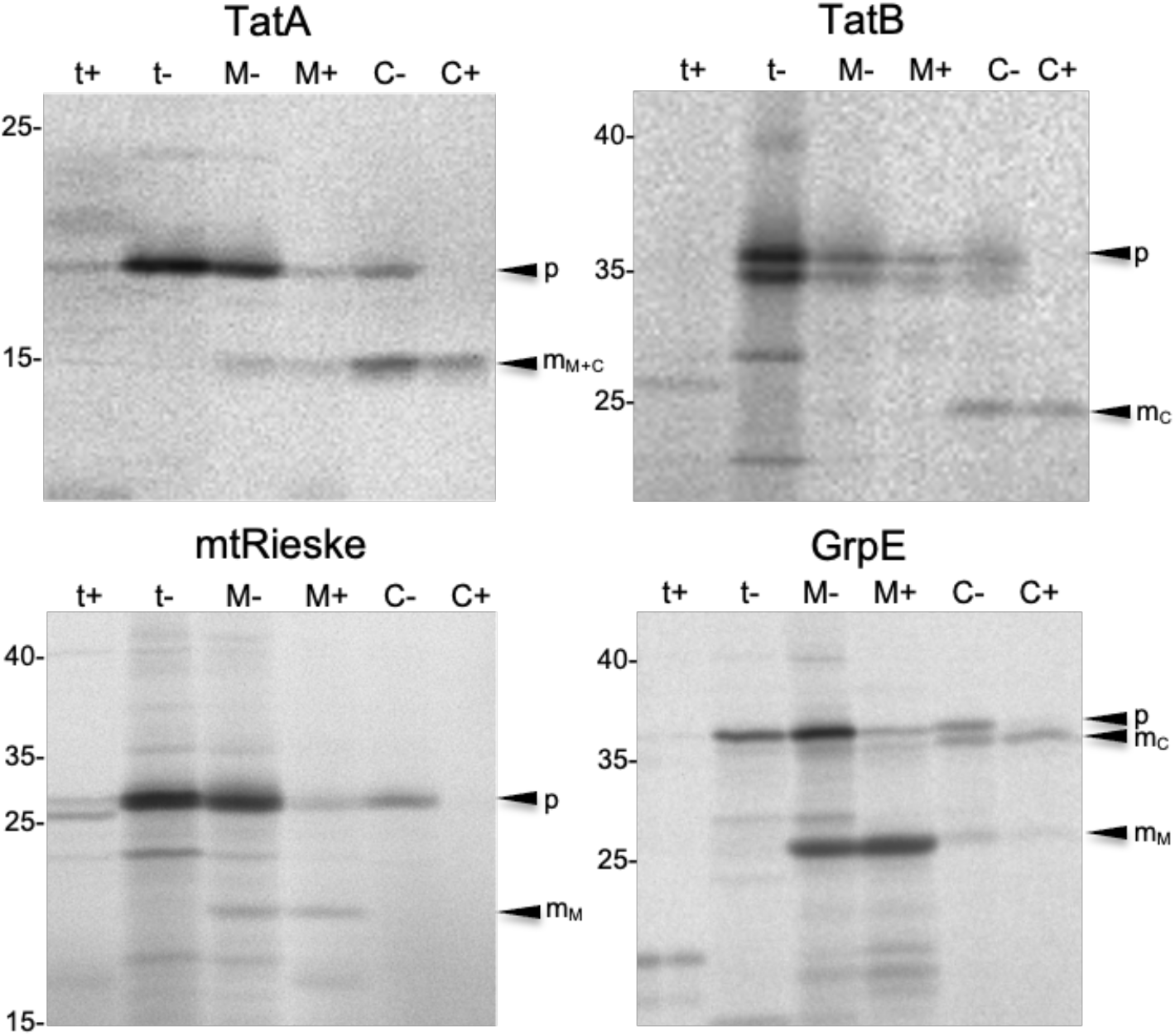
*In organello* protein transport experiments with isolated pea organelles. Radiolabelled precursor polypeptides of peaTatA (*TatA*) and the control proteins chloroplast TatB (*TatB*), mitochondrial Rieske Fe/S protein (*mtRi*), and the dually targeted GrpE (*GrpE*) were obtained by coupled *in vitro* transcription/translation of the corresponding cDNA clones in the presence of [^35^S]-methionin and incubated for 20 min at 25 °C with either intact mitochondria (*lanes M*) or chloroplasts (*lanes C*) from pea. After the import reaction, the organelles were recovered by centrifugation and treated with thermolysin (*lanes M+, C+*) or mock-treated (*lanes M−, C−*). Stoichiometric amounts of each fraction corresponding to 50 μg protein (mitochondria) or 12.5 μg chlorophyll (chloroplasts) were separated on 10 −17.5 % SDS polyacrylamide gradient gels and visualised by phosphorimaging. In *lanes t*, 1 μl aliquots of the *in vitro* translation assays (corresponding to 10 % of the protein added to each import reaction) were loaded that had been either treated with thermolysin (*lanes t+*) or mock-treated (*lanes t−*). The positions of the precursor proteins (p) and the mature proteins accumulating in mitochondria (*m_M_)* or chloroplasts (*m_C_*) are indicated. The size of molecular marker proteins is given in kDa.

### In organello competition experiments

Such differential processing within the two organelles, as for GrpE, is not observed for TatA though. Instead, both in mitochondria and chloroplasts the precursor is cleaved to terminal processing products of apparently identical size, as was observed earlier also for other dually targeted proteins (e.g. Baudisch and Klösgen, 2012; Baudisch et al., 2014). This lack of distinctiveness makes it more difficult to rule out that the observed import into mitochondria is not in fact caused by contaminating plastids.

One way to distinguish between chloroplast and mitochondrial import of a candidate protein are competition experiments in which saturating amounts of an unlabelled precursor protein with known organelle targeting specificity are added together with the radiolabelled candidate protein to the *in organello* assays (Rödiger et al., 2011; Baudisch and Klösgen, 2012; Langner et al., 2014). Here we have used the precursor of the mitochondrial Rieske protein as competitor, which was obtained by heterologous overexpression in *E. coli* and subsequent purification. In the presence of increasing amounts of this mitochondria-specific competitor, the accumulation of mature TatA within mitochondria was successively reduced, whereas its accumulation in chloroplasts remained unaffected (Fig. 5A,B). Instead, if the chloroplast-specific OEC33 competitor was added in increasing amounts to the assays, import of TatA into chloroplasts was successively reduced (Fig. 5C and suppl. Fig. S3A). Unexpectedly, import of TatA into mitochondria was also affected to some extent in the presence of the OEC33 competitor but not in a concentration-dependent manner. At all competitor concentrations ranging from 0.2 - 3.2 μM in the assays, mitochondrial import of TatA is dropped to 40 - 60% (Fig. 5C). At first glance, this might suggest competition of import into contaminating plastids in the assays. However, the same behavior was observed also when the transport of the monospecific mitochondrial Rieske protein into mitochondria was analyzed (suppl. Fig. S3B). Despite the fact that mtRieske cannot at all be imported into chloroplasts (suppl. Fig. S3B and Baudisch et al., 2014), import of mtRieske in the mitochondrial assays is dropped to 40% in the presence of the OEC33 competitor and, again, there is no clear concentration-dependent effect visible (Fig. 5E). The reason for this impairment remains unclear yet. Still, these data demonstrate that the observed accumulation of mature TatA within mitochondria is the result of true mitochondrial import and not caused by the presence of contaminating plastids in the assays.

**Figure 5.**
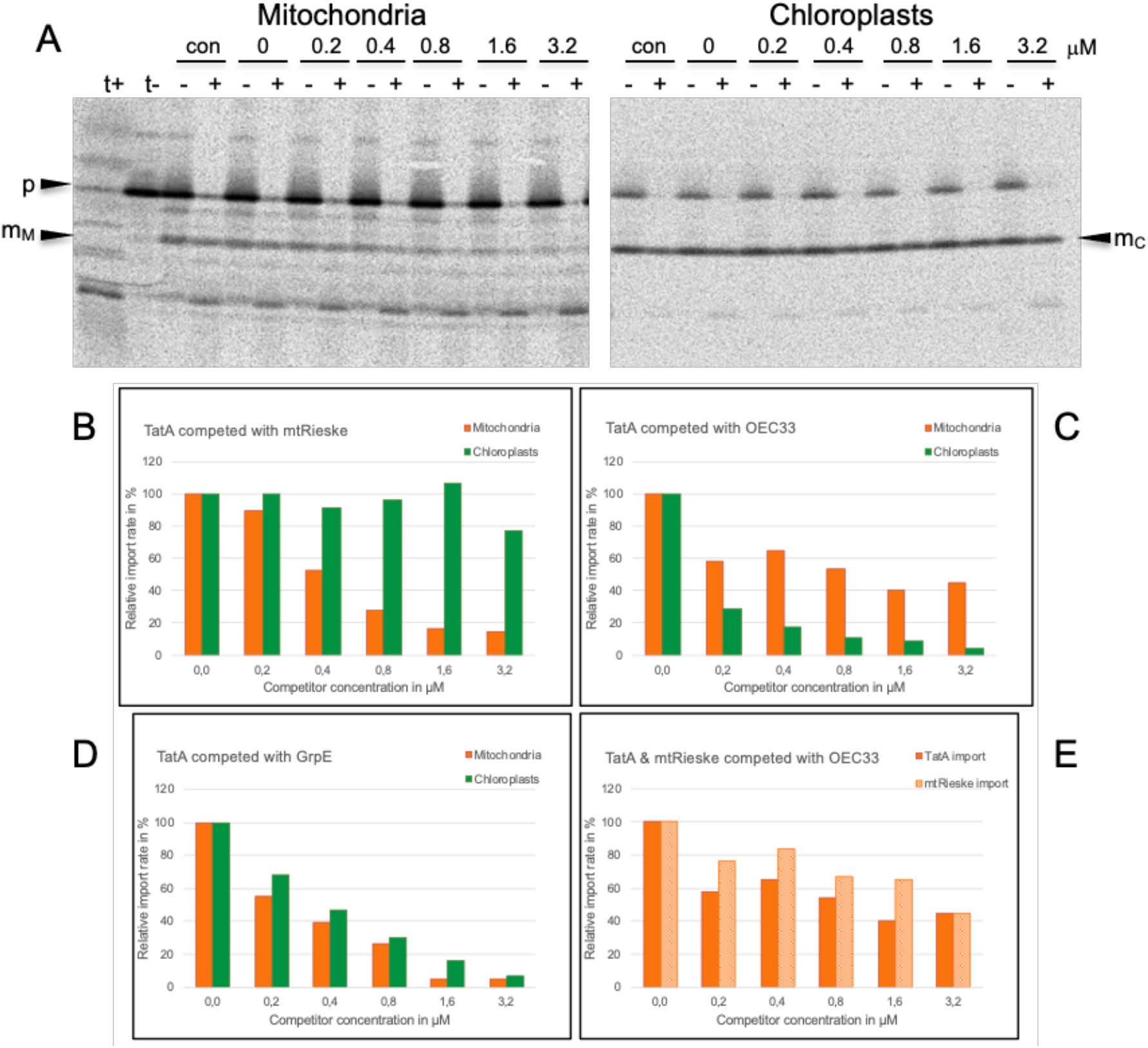
Effect of competitor proteins on organelle import of TatA. **(A)** *In organello* protein transport experiments with the TatA precursor protein were performed in the absence (*con*) or presence of increasing amounts of the precursor of the mitochondrial Rieske Fe/S protein which was obtained by heterologous overexpression in *E. coli*. The concentration of competitor protein present in each assay (given in μM) is indicated above the lanes. The bands corresponding to mature TatA in *lanes* + were quantified and depicted in terms of percentage of mature TatA in the assays lacking competitor (*lanes 0*) in the bar graph shown in **(B)**. Panels **(C)** and **(D)** show the same experiment performed in the presence of the chloroplast-specific competitor protein OEC33 **(C)** and the dually targeted competitor protein GrpE **(D)**, respectively. In panel **(E),** the effect of the competition with the OEC33 competitor on the mitochondrial import of TatA and mtRieske is compared. The autoradiograms of the gels used for the quantification data shown in panels **(C-E)** are presented in suppl. Fig. S3. For further details see the legend of Fig. 4.

This conclusion was finally confirmed by complementing experiments employing GrpE as competitor to saturate organelle import of TatA. In this case, competition of TatA import into both organelles, mitochondria and chloroplasts, was observed (suppl. Fig. S3C), in line with the dual targeting properties of GrpE (Fig. 4 and Baudisch et al., 2014). Even more, the competition curves obtained with the GrpE competitor look almost identical for both organelles (Fig. 5D) indicating that TatA and GrpE show a similar affinity pattern for each organelle.

## Discussion

It was the goal of this study to examine whether the TatA component of the thylakoidal Tat machinery, which is encoded in the nucleus of the cell, is imported also into mitochondria of plant cells. If so, it would suggest that TatA is not only active in the thylakoidal Tat pathway but might also serve as a constituent subunit of the recently identified plant mitochondrial Tat machinery. As yet, this machinery is assumed to consist of a mitochondrially encoded TatC-orthologue (OrfX, Sünkel et al., 1994) and a nuclearly encoded TatB-like protein, a structural analogue of thylakoidal TatB (Carrie et al., 2016). The latter was recently shown to be essential for the biogenesis of complex III of the respiratory electron transport chain, which strongly suggests that a Tat-like transport pathway is operating also in plant mitochondria (Schäfer et al., 2020).

### The TatA precursor is recognized by the import machineries of both chloroplasts and mitochondria

The major outcome of this study is that TatA is indeed dually targeted into both, mitochondria and chloroplasts. As shown with *in organello* experiments (Fig. 4), incubation of the TatA precursor with either of the two endosymbiotic organelles leads to the organellar accumulation of a processing product corresponding in size to mature chloroplast TatA. Import of the protein into mitochondria proceeds by the general import pathway for nuclearly encoded proteins of the mitochondrial matrix as was confirmed by *in organello* competition experiments employing the precursor of the mitochondrial Rieske protein as competitor (Fig. 5).

However, such mitochondrial import of TatA cannot easily be detected with standard *in vivo* assays resting on fluorescent reporter constructs like eYFP. In the biolistic transformation experiments performed here, mitochondrial targeting of the TatA/eYFP chimera was found in only 5 - 10% of the transformed cells, while in the majority of cases solely import into chloroplasts could be detected (Fig. 1A). After agroinfiltration of *Nicotiana benthamiana* leaves, import of the same reporter construct into mitochondria could not even be discovered at all (Fig. 2A). This seeming lack of mitochondrial import is in fact a matter of too low sensitivity of such assays. When the agroinfiltration experiments were performed with the self-assembling split green fluorescent protein (*sa*split-GFP) as reporter (Cabantous et al., 2005), transport of TatA into mitochondria of *Nicotiana* leaves was clearly visible (Fig. 3B), presumably due to the lower background fluorescence in these assays. Hence, TatA is dually targeted into both, chloroplasts and mitochondria, but mitochondrial import takes place with such low rates in the tissues analyzed here that it can be discovered only with highly sensitive detection methods.

### Biological relevance of mitochondrial Tat transport

The apparent lower efficiency of TatA transport into mitochondria vs. chloroplasts might point to a low demand for Tat translocase in this organelle. To date, solely a single substrate of this pathway has been identified which is the mitochondrial Rieske Fe/S protein (Schäfer et al., 2020). Further substrates might be detected in the future but it appears unlikely that the number will raise very much considering that in bacteria Tat transport is largely restricted to proteins that demand for cytoplasmic insertion of metal ions or complex cofactors prior to membrane translocation (Palmer and Berks, 2012), like NADPH (Halbig et al., 1999) or Fe/S clusters (Molik et al., 2001). In chloroplasts, Tat transport is more common and comprises substrate proteins with (cpRieske Fe/S protein) and without such cofactors (OEC16, OEC23) (Klösgen et al., 1992; Cline et al., 1992; Molik et al., 2001) maybe because the higher energy demand for the membrane transport of folded proteins (Alder and Theg, 2003) is not that decisive in light-exposed chloroplasts.

### Components of the plant mitochondrial Tat machinery

Our results on the transport characteristics of TatA suggest that in plant mitochondria a chloroplast-like Tat pathway is operating. This means that the mitochondrial Tat machinery probably comprises all three Tat subunits identified to date, namely TatA, a TaB-like protein, and mitochondrial TatC (OrfX), in accordance with Gram-negative bacteria and chloroplasts. Differently composed Tat translocases are found in most Gram-positive bacteria including *Bacillus subtilis* which houses a Tat machinery consisting of only two subunits, namely TatA and TatC (Jongbloed et al., 2004). In this case, TatA has a bifunctional role, i.e., it provides both, TatA and TatB activity (Blaudeck et al., 2005; Barnett et al., 2008; Jongbloed et al., 2006). A comparable, bifunctional role was recently suggested also for the mitochondrial TatB-like protein to explain the presumed lack of a TatA component in mitochondria (Schäfer et al., 2020). This cannot finally be excluded at this point but appears less likely, considering the import of TatA into mitochondria observed here. Furthermore, a mitochondrial Tat translocase of the TatABC-type would be in accordance also with the evolutionary origin of this endosymbiotic organelle from Gram-negative alpha-proteobacteria (Gray et al., 1999).

In this scenario, it is particularly remarkable that the Tat machineries of the two plant organelles would use one subunit in common, in spite of the fact that the two other Tat components in chloroplasts and mitochondria (TatB and TatB-like as well as nuclearly encoded chloroplast TatC and mitochondrially encoded TatC/OrfX, respectively) are in each case quite different from each other. The two TatC proteins show only about 20 - 25% amino acid identity to each other, while TatB and the TatB-like protein have even entirely different sequences despite the fact that they possess an essentially identical structure (Carrie et al., 2016). Yet, if the dually targeted TatA protein indeed plays an active role also in mitochondrial Tat transport, which still remains to be proven though, it would add one further peculiarity to this already enigmatic membrane transport pathway.

## Methods

### Plant material

Pea seedlings (*Pisum sativum* var. Feltham First) were grown on soil for 7 - 10 days at a 16 h photoperiod under constant temperature (18-22 °C). *Nicotiana benthamiana* was grown at the same conditions for 6 - 8 weeks.

### Molecular Cloning

The cDNA fragments encoding the precursor proteins of TatA and TatB from pea (gene accession numbers AF144708 and AF284760, respectively) were amplified by PCR using a cDNA library generated from pea seedlings as template (see suppl. Table 1 for all primer sequences). The amplified fragments were cloned with the pBAT vector (Annweiler et al., 1991) and subsequently modified to introduce methionine codons to permit labeling of the mature proteins with [35S]methionine. In TatA, the methionine codon was inserted immediately upstream of the stop codon, while in TatB the valine codon at position 245 of the precursor protein was substituted. Mutagenesis was carried out with the QuikChange^®^ Site-Directed Mutagenesis Kit (Stratagene, La Jolla, CA, USA) and confirmed by DNA sequencing.

Cloning of the cDNAs encoding the precursors of mitochondrial Rieske Fe/S protein from potato and the nucleotide exchange factor GrpE from *Arabidopsis thaliana* are described in Emmermann et al. (1994) and Baudisch et al. (2014), respectively. For bacterial overexpression of the two precursor proteins, the cDNAs were amplified by PCR introducing NdeI restriction sites at the start codons and recloned with vector pET30a (Novagen) yielding chimeric fusion constructs with C-terminal hexahistidine-tags.

The constructs used for biolistic transformation of pea epidermal cells are based on the vector pRT100mod (pRT100 Ω/Not/Asc; Überlacker and Werr, 1996) and have been described in Baudisch et al. (2014). For the generation of the peaTatA_1-100_/eYFP and AtTatB_1-100_/eYFP constructions, cDNA fragments encoding the N-terminal 100 residues were amplified via PCR and cloned in frame with the coding sequence of the enhanced yellow fluorescent protein (eYFP) via restriction digestion and subsequent ligation. The subsequent cloning with the binary vector pCB302 (Xiang et al., 1999) needed for *Agrobacterium* infiltration is described in Sharma et al. (2018).

Likewise, the generation of the *sa*split/GFP reporter constructs, namely FNR_1-55_/GFP1-10, FNR_1-55_/GFP11_x7_, mtRi_1-100_/GFP1-10, and mtRi_1-100_/GFP11_x7_, are described in Sharma et al. (2019). The coding sequence of eYFP in pRT100peaTatA_1-100_/eYFP was replaced by GFP11_x7_ via restriction-free cloning (Bond and Naus, 2012) generating pRT100peaTatA_1-100_/GFP11_x7_. Subsequently, the peaTatA_1-100_/GFP11_x7_ fragment was sub-cloned with the binary vector pLSU4GG using Golden Gate cloning as described in Sharma et al. (2018).

### Bacterial overexpression and purification of precursor proteins

His-tagged precursor proteins of mtRieske from potato and GrpE from *Arabidopsis thaliana* were obtained by overexpression in *E. coli* strain BL21(DE3) using the T7-based system developed by Studier and Moffat (1986) following the protocol of Zinecker et al. (2020) with the following modifications: (i) cultures were grown in LB media supplemented with 0.4% glucose, 50 mg/ml kanamycin for 1.5 - 3 h after induction with IPTG, (ii) the His-tagged proteins were recovered from inclusion bodies solubilized in GuaHCl binding buffer (20 mM Hepes, 500 mM NaCl, 20 mM Imidazole, 6 M GuanidinHCl, pH 7.5), (iii) during Ni^2+^-affinity chromatography the column (HiTrap Chelating HP columns, GE Healthcare) was washed with (20 mM Hepes, 500 mM NaCl, 20 mM Imidazole, 8 M urea, pH 7.5) and eluted with (20 mM Hepes, 500 mM NaCl, 500 mM Imidazole, 8 M urea, pH 7.5), (iv) the samples were concentrated using Vivaspin 3.000 MWCO PES ultrafiltration columns (Sartorius AG, Göttingen, Germany) and finally dialysed against (10 mM Hepes, 5 mM MgCl_2_, 7 M urea, pH 8.0).

### Transient transformation assays

Particle bombardment of 7 - 10 days old pea seedlings was carried out according to Sharma et al. (2018). *Agrobacterium* infiltration of eYFP reporter constructs and sasplitGFP vectors into the lower epidermis of *Nicotiana benthamiana* leaves were performed as described (Sharma et al., 2018 and 2019, respectively). The mitochondria of epidermal cells were stained by infiltrating MitoTracker Orange (0.1 μM MitoTracker Orange® CMTMRos in 10 mM MgCl_2_, 10 mM MES, pH 5.6) into the lower epidermis of leaf tissue and imaged 15 min after infiltration.

### Microscopy and imaging

Confocal laser scanning microscopy was carried out as described by Sharma et al. (2018). The specimens were excited with laser beams of either 488 nm (*sa*split-GFP), 514 nm (eYFP), 561 (MitoTracker Orange), or 633 nm (chlorophyll) with usually 2-5% of full laser power. Images were collected using filters ranging from 493 to 598 nm (GFP), 519–620 nm (eYFP), 574-617 nm (MitoTracker Orange), or 647–721 nm (chlorophyll). Image acquisition was done in several Z-stacks with either 20x or 40x magnification. When required, brightness and contrast of the images were equally adjusted for each image to avoid any discrepancy in visualization of signal intensities.

### Isolation of chloroplasts and mitochondria from pea

Isolation of intact chloroplasts and mitochondria from pea seedlings (*P. sativum* var. Feltham First) was carried out according to Rödiger et al (2010) with the following modifications: The plant material was homogenized in sucrose isolation medium (SIM: 0.35 M sucrose, 25 mM Hepes, 2 mM EDTA, pH 7.6) supplemented with 0.6% PVP, 0.2% BSA, 10mM DTT, 0.2 mM PMSF. The Percoll step gradients for isolation of mitochondria consisted of 5 ml each of 40%, 30%, and 18.5% Percoll in sorbitol resuspension medium (SRM: 0.35 M sorbitol, 50 mM Hepes, pH 8.0).

### In organello protein transport experiments

Radiolabelled precursor proteins were obtained by *in vitro* translation in rabbit reticulocyte lysates in the presence of [^35^S]-methionine. Incubation with intact mitochondria or chloroplasts isolated from pea leaves was performed as described in Rödiger et al. (2010) with the exception that the transport assays were additionally supplemented with 2 mM NADH. Competition experiments were performed as described (Molik et al., 2001; Langner et al, 2014).

### Miscellaneous

Gel electrophoresis of proteins under denaturing conditions was carried out according to Laemmli (1970). The gels were exposed to phosphorimaging screens and analyzed with a Fujifilm FLA-3000 (Fujifilm, Düsseldorf, Germany) using the software packages BAS Reader (version 3.14) and AIDA (version 3.25) (Raytest, Straubenhardt, Germany). Protein concentration was determined according to Bradford (1976), chlorophyll concentration according to Arnon (1949). All other methods followed published protocols (Sambrook & Russell, 2001).

## Abbreviations

Tat: Twin-arginine translocation
OEC33: 33 kDa subunit of the oxygen-evolving system associated with photosystem II
sfGFP: ‘superfolder’ GFP
Hepes: 4-(2-Hydroxyethyl)piperazine-1-ethanesulfonic acid
PVP: Polyvinylpyrrolidone
PMSF: Phenylmethylsulfonyl fluoride

## Author Contributions

BB performed the *in organello* experiments. MS performed the *in vivo* experiments. FT isolated and purified the precursor proteins for competition. RBK designed and supervised the project. BB, MS, and RBK wrote the manuscript.

## Acknowledgements

This work was supported by a grant from the Deutsche Forschungsgemeinschaft (GRK2498 - 400681449). MS was supported by a fellowship from the BRAVE project funded by the ERASMUS MUNDUS Action 2 program of the European Union. We thank Mario Jakob for critical comments.

## Supplemental Information

**Supplemental Figure S1.**
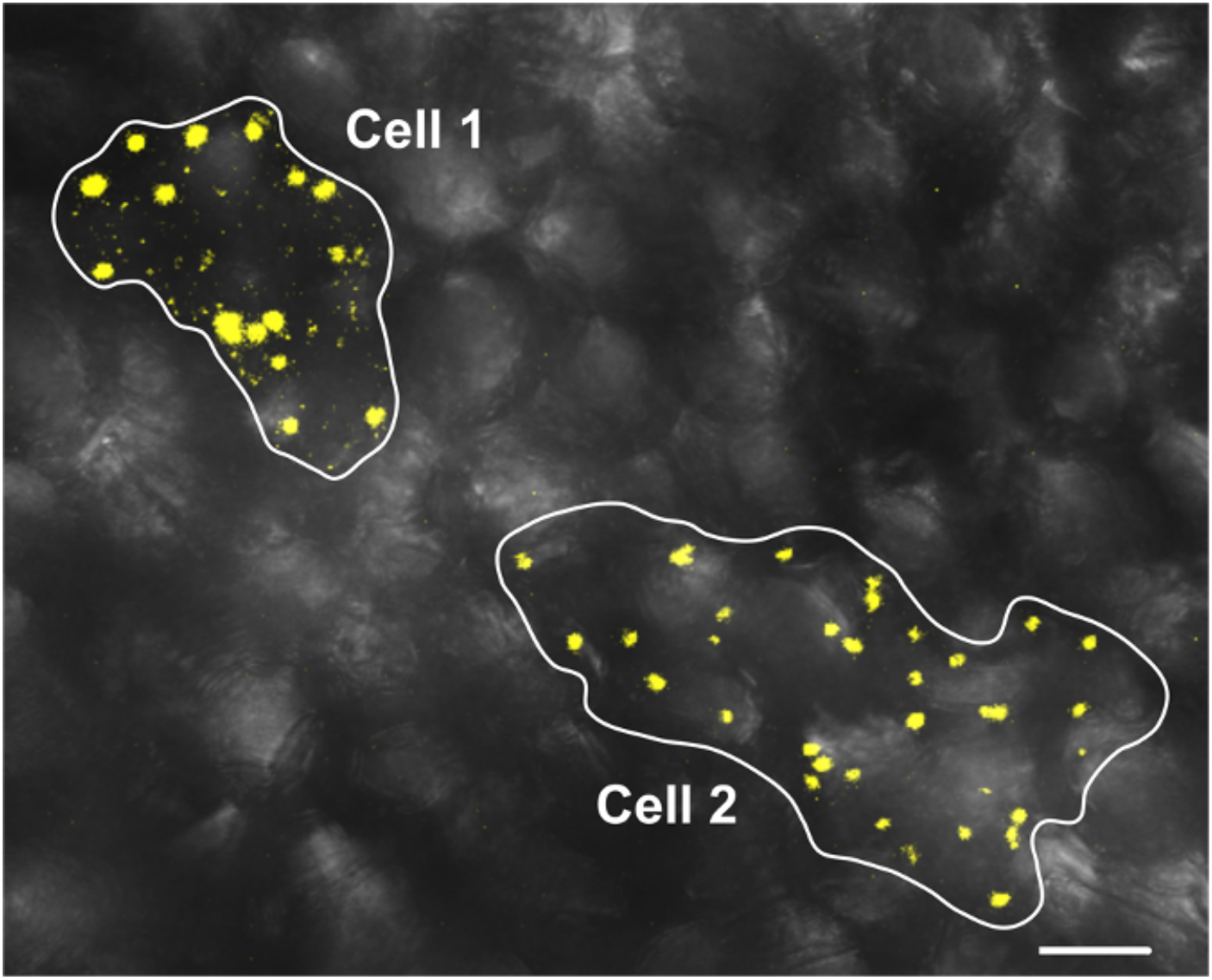
Differential localization of the TatA_1-100_/eYFP chimera in two neighboring pea cells after particle bombardment. The coding sequence of peaTatA_1-100_/eYFP was transiently expressed under the control of the CaMV 35S promoter after particle bombardment of leaf epidermis cells of pea and analyzed by confocal laser scanning microscopy. Cell 1 shows dual localization of eYFP in plastids and mitochondria, while Cell 2 shows localization of eYFP in plastids only. The image is presented as an overlay of DIC (differential interference contrast) and eYFP channels. For further details see the legend of Fig. 1. Scale bar: 20 μm.

**Supplemental Figure S2.**
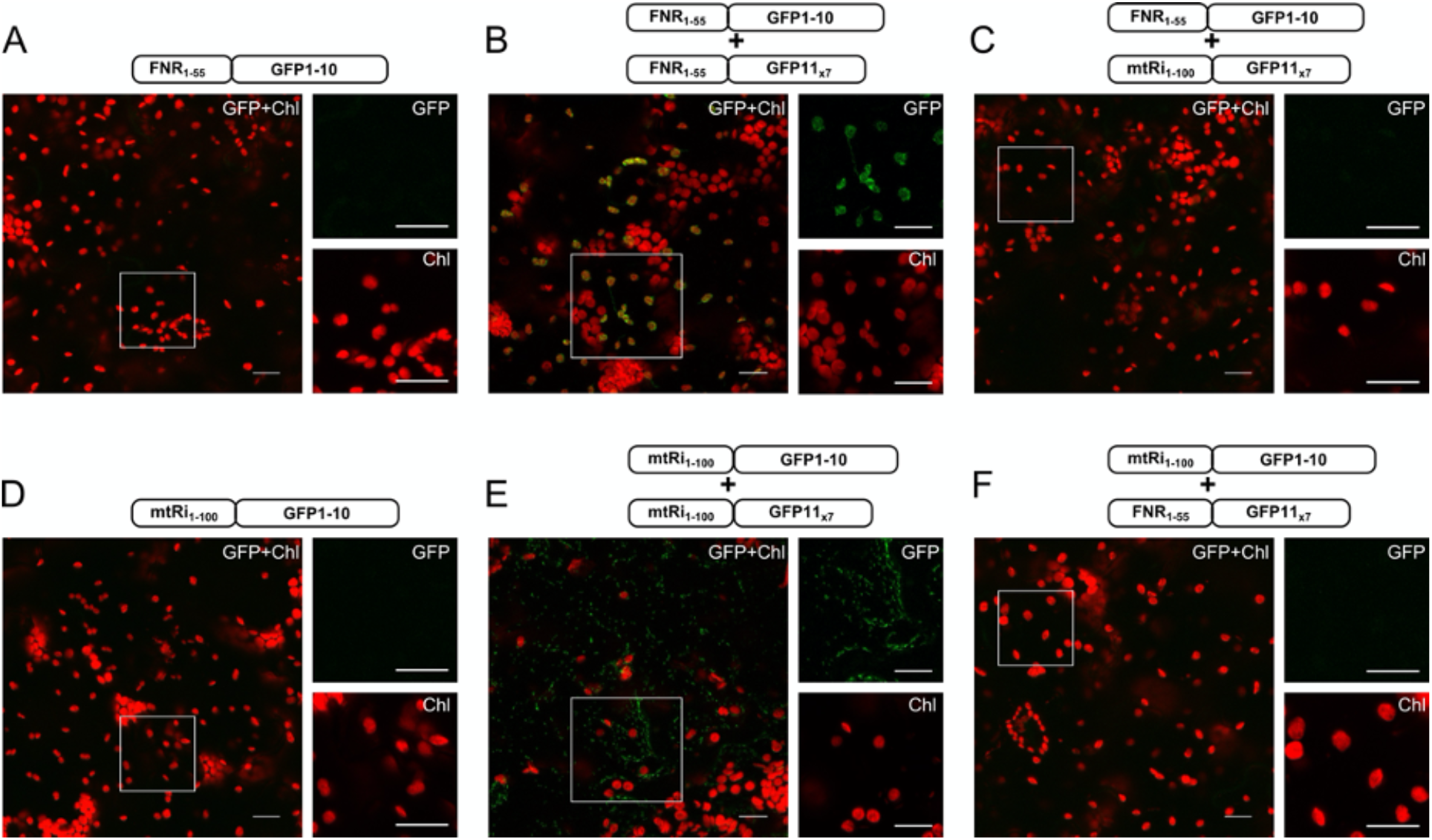
Transport of both fragments of sfGFP into the same organelle is essential to achieve fluorescence signals. The coding sequences of FNR_1-55_/GFP1-10 **(A-C)** or mtRi_1-100_/GFP1-10 **(D-F)** were transiently expressed alone **(A,D)** or co-expressed with either FNR_1-55_/GFP11_×7_ **(B,F)** or mtRi_1-100_/GFP11_x7_ **(C,E)** via *Agrobacterium* co-infiltration into the lower epidermis of *Nicotiana benthamiana* leaves and analyzed by confocal laser scanning microscopy. For further details see the legend of Fig. 1. Scale bars: 20 μm.

**Supplemental Figure S3.**
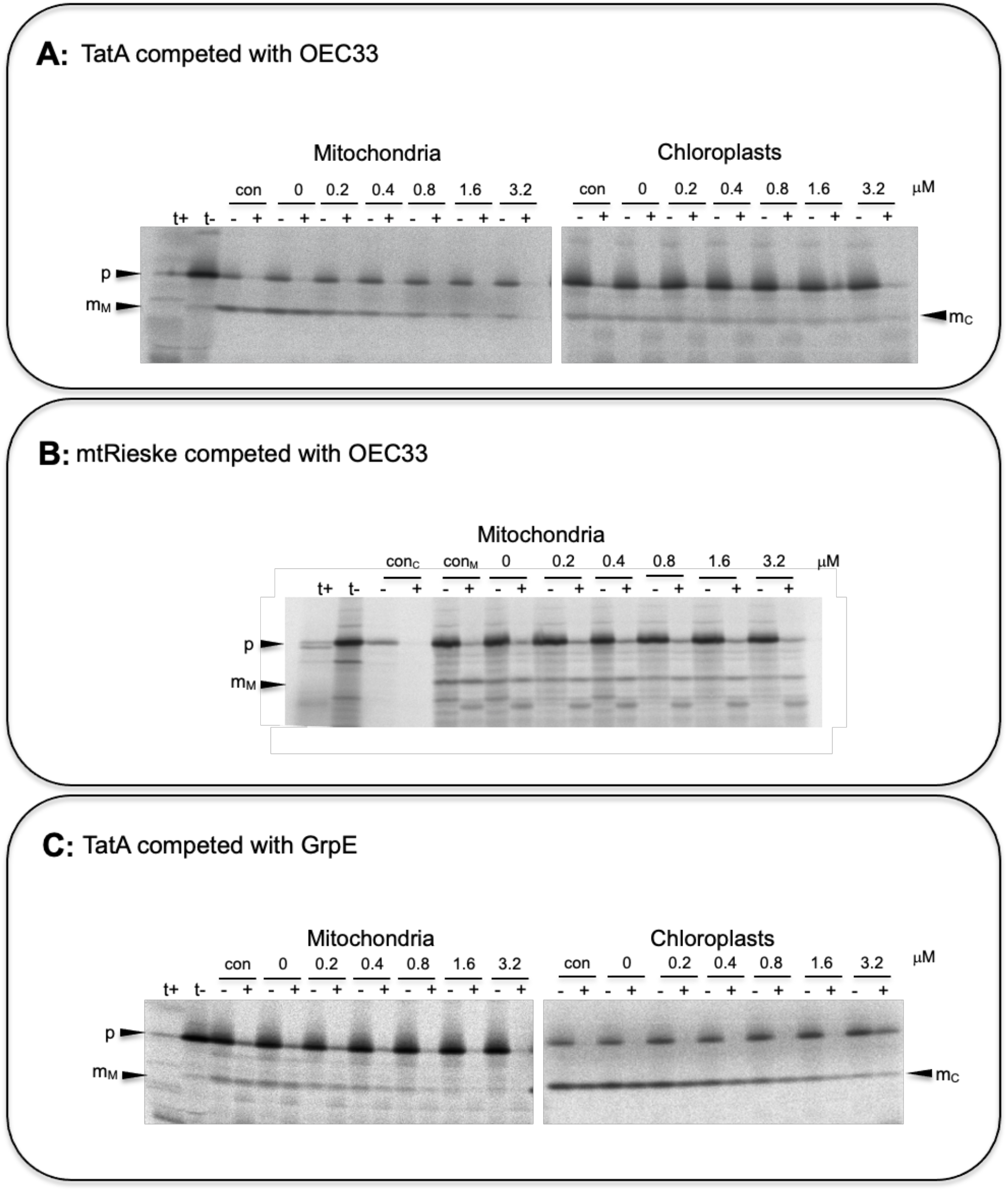
*In organello* competition experiments. *In organello* protein transport experiments with **(A)** the TatA precursor protein, or **(B)** the mtRieske precursor performed in the absence (*con*) or presence of increasing amounts of the chloroplast-specific competitor OEC33. In panel **(C)**, organelle transport of TatA in the presence of the dually targeted competitor GrpE is shown. For further details see the legends of Figs. 4 and 5.

**Supplementary Table 1.**
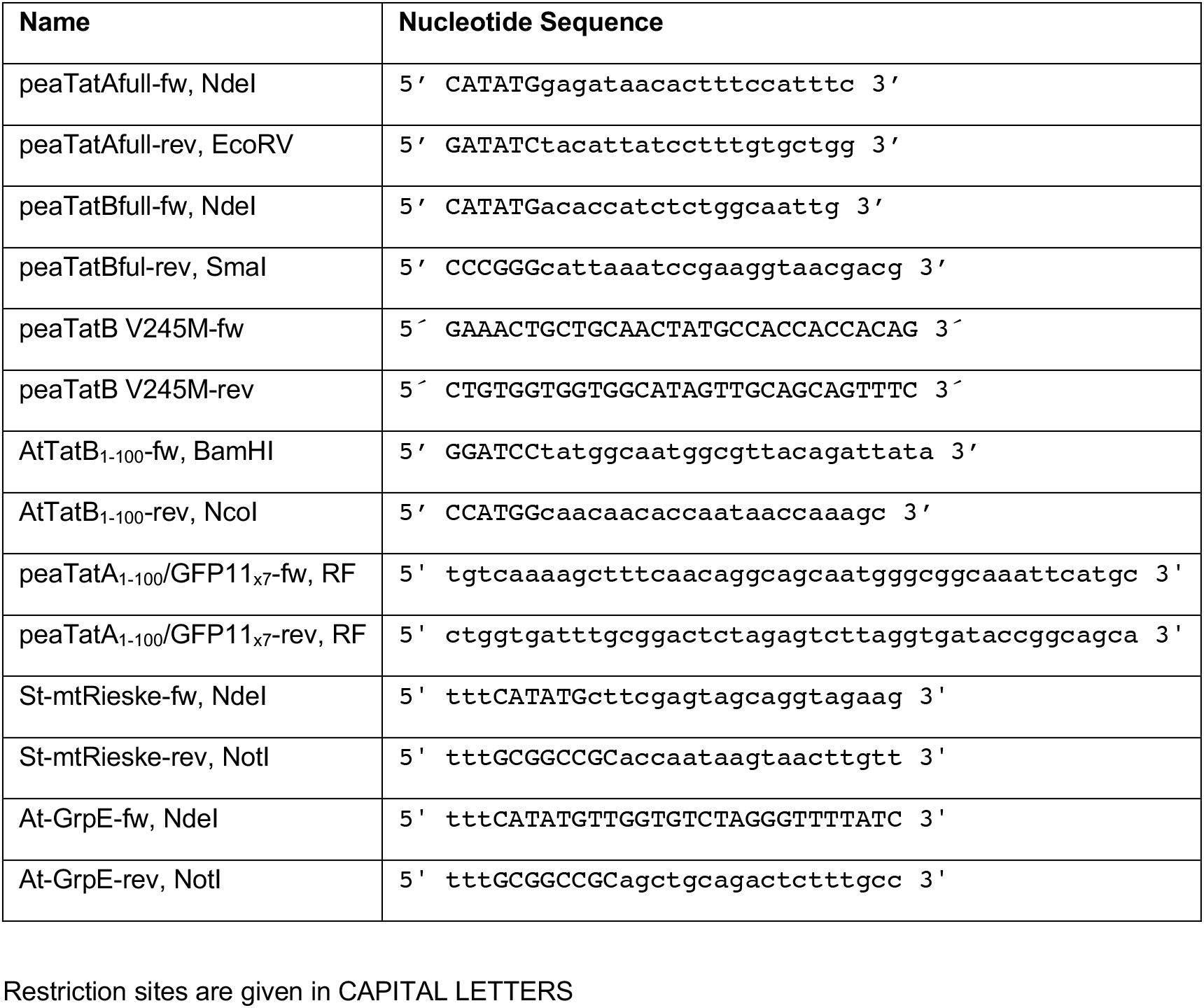
Oligonucleotides.

